# Gene expression shapes the patterns of parallel evolution of herbicide resistance in the agricultural weed *Monochoria vaginalis*

**DOI:** 10.1101/2021.03.10.434542

**Authors:** Shinji Tanigaki, Akira Uchino, Shigenori Okawa, Chikako Miura, Kenshiro Hamamura, Mitsuhiro Matsuo, Namiko Yoshino, Naoya Ueno, Yusuke Toyama, Naoya Fukumi, Eiji Kijima, Taro Masuda, Yoshiko Shimono, Tohru Tominaga, Satoshi Iwakami

## Abstract

The evolution of herbicide resistance in weeds is an example of parallel evolution, through which genes encoding herbicide target proteins are repeatedly represented as evolutionary targets. The number of herbicide target-site genes differs among species, and little is known regarding the effects of duplicate gene copies on the evolution of herbicide resistance. We investigated the evolution of herbicide resistance in *Monochoria vaginalis*, which carries five copies of sulfonylurea target-site acetolactate synthase (*ALS*) genes. Suspected resistant populations collected across Japan were investigated for herbicide sensitivity and *ALS* gene sequences, followed by functional characterisation and *ALS* gene expression analysis. We identified over 60 resistant populations, all of which carried resistance-conferring amino acid substitutions exclusively in *MvALS1* or *MvALS3*. All *MvALS4* alleles carried a loss-of-function mutation. Although the enzymatic properties of ALS encoded by these genes were not markedly different, the expression of *MvALS1* and *MvALS3* was prominently higher among all *ALS* genes. The higher expression of *MvALS1* and *MvALS3* is the driving force of the biased representation of genes during the evolution of herbicide resistance in *M. vaginalis*. Our findings highlight that gene expression is a key factor in creating evolutionary hotspots.

## INTRODUCTION

Agricultural weeds are often subject to artificial lethal stresses employed in an effort to increase crop yield. Such weeding stresses, in turn, act as selection pressures driving the evolution of weed populations. While multiple evolved traits related to weed adaptation in agricultural ecosystems have been documented, such as flowering phenology, mimicry morphology, seed shattering, seed dormancy, and competitiveness (Fukano, Guo, Uchida, Tachiki, & Cornelissen, 2020; Waselkov, Regenold, Lum, & Olsen, 2020; Wedger & Olsen, 2018; Ye et al., 2019), herbicide resistance remains among the best studied and characterised trait (Powles & Yu, 2010). Evolved herbicide resistance has been documented worldwide in as many as 263 weed species (Heap, 2021), posing a great threat to current global food security.

While weed resistance has often been studied from an agricultural viewpoint as a threat to food security, it has also attracted the attention of evolutionary biologists as a classic example of an ‘evolution in action’ (Baucom, 2019). An important insight from studies on weed resistance is that nature often mimics itself: certain genes have repeatedly acted as the major components in the evolution of resistance (Martin & Orgogozo, 2013). Research so far has observed that weed populations subjected to herbicide applications evolved resistance via an identical amino acid substitution (AAS) in the genes encoding herbicide target sites, irrespective of evolutionary lineage (Baucom, 2016). These mutations result in conformational alterations in the binding site of each herbicide, leading to the herbicide-insensitive form of the protein (Powles & Yu, 2010). Although this parallelism has been observed in many herbicides with diverse modes of action, resistance to herbicides inhibiting acetolactate synthase (ALS, or acetohydroxy acid synthase), in particular, represents a case of extreme parallelism at the molecular level (Yu & Powles, 2014). Due to their low toxicity to animals, high crop selectivity, and high efficacy at low doses, ALS herbicides are among the most frequently used herbicides in many countries, including Japan (Hamamura, 2018; Peters & Strek, 2018). The ALS residues where AASs induce resistance have been reported in as many as eight from natural weed populations, and most of these mutations were not significantly associated with fitness cost (Yu & Powles, 2014). This nature is considered as the driving force to render *ALS* loci as an evolutionary hotspot for resistance to ALS herbicides (Baucom, 2019).

Little attention has been paid to the effects of duplicated gene copies on the evolution of herbicide resistance. Having undergone whole genome duplications during the long course of evolution (Clark & Donoghue, 2018), many plants possess duplicated copies of target-site genes even in diploid species, e.g. *ALS* genes in *Senecio vulgaris* and *Zea mays*(Délye, Causse, & Michel, 2016; Svitashev et al., 2015) and acetyl-CoA carboxylase genes in *Echinochloa* spp. (Iwakami et al., 2015; Iwakami, Uchino, Watanabe, Yamasue, & Inamura, 2012). Genetic redundancy by duplicated copies fosters the evolutionary differentiation of each copy, such as gene expression and/or function of the encoded protein, including pseudogenisation (Flagel & Wendel, 2009). Although not much is understood in the differentiation of herbicide target-site genes in weeds, the differentiation, if any, may provide distinct competency for resistance evolution in each “potentially hot” locus, influencing the evolutionary dynamics of resistance mechanisms.

*Monochoria vaginalis* (syn. *Pontederia vaginalis*) is a noxious allohexaploid (2n = 4x = 52) weed in temperate and tropical Asian paddy fields (Holm, Plucknett, Pancho, & Herberger, 1977). It carries five *ALS* loci, with all genes transcribed at the seedling stage (Iwakami et al., 2020) when plants are exposed to herbicides in agricultural fields. Among the limited number of populations tested to date, mutations known to confer ALS herbicide resistance have been reported exclusively in *MvALS1* or *MvALS3* (e.g. Iwakami et al., 2020; Ohsako & Tominaga, 2007), despite the presence of multiple potential evolutionary hotspots for resistance. In the present study, we confirmed the occurrence of resistance-conferring mutations exclusively in *MvALS1* and *MvALS3* among 68 resistant populations across Japan. Enzyme kinetics and expression analysis of each *ALS* copy and its product unveiled the potential mechanisms that make these two loci the hotspots of resistance evolution.

## MATERIALS AND METHODS

### M. vaginalis

*M. vaginalis* was collected from paddy fields across Japan (Figure 1a-b; Table S1). Most of the paddy fields were small, measuring 0.1 to 0.3 ha an average. Plants from a single paddy field are hereafter referred to as a population. Adjacent fields were avoided during sampling. The plants were either directly used for resistance diagnosis using the rooting assay or cultivated for the collection of selfed seeds for the following pot assay (Figure 1c). The phenotyped plants were subjected to *ALS* gene sequencing (see section 2.4).

**Figure 1.**
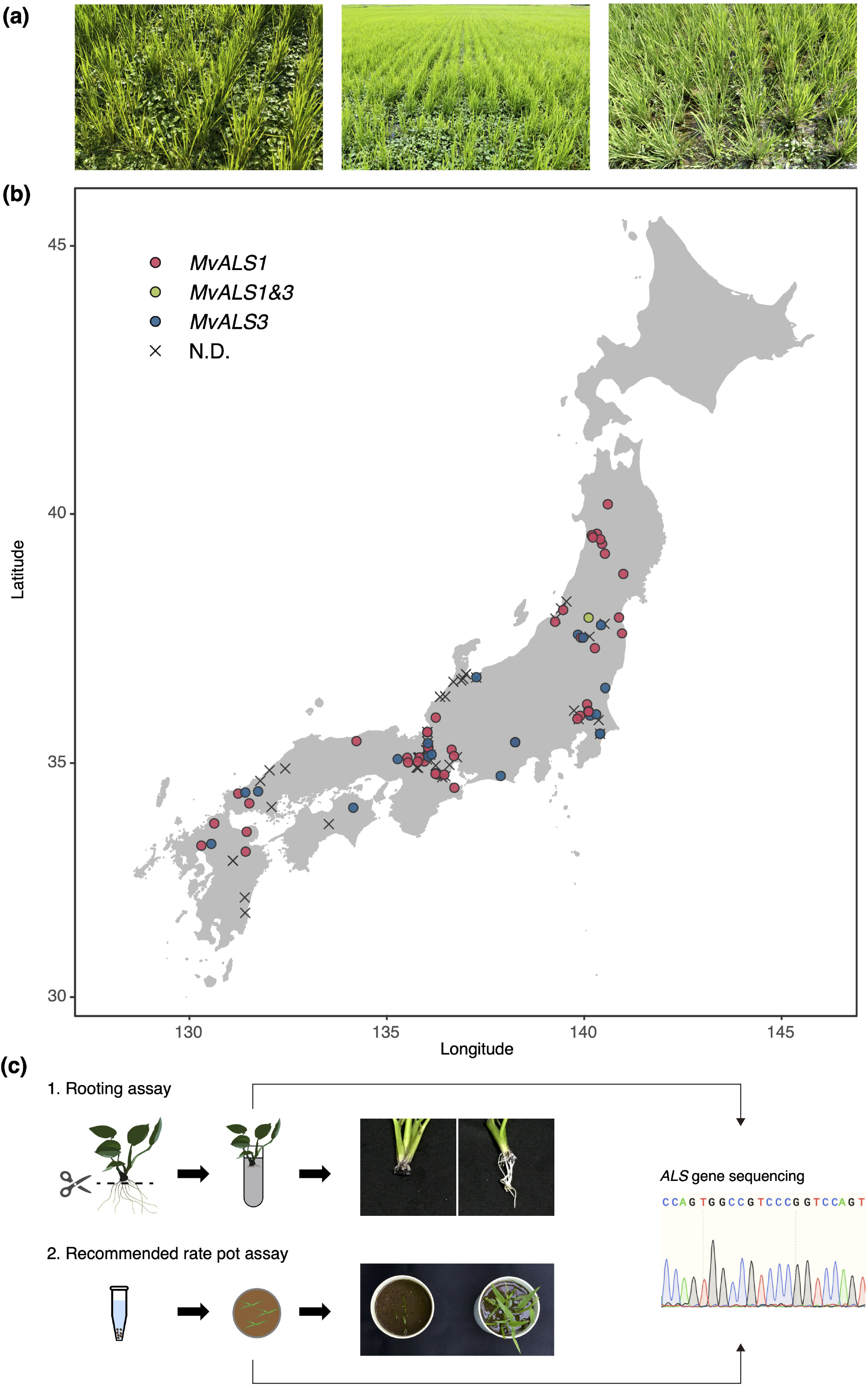
*Monochoria vaginalis* studied in this work. (a) Infestation of *M. vaginalis* in paddy fields in Japan. (b) Collection sites of *M. vaginalis*. *MvALS1*, populations carrying resistance-conferring mutations in *MvALS1; MvALS3*, populations carrying resistance-conferring mutations in *MvALS3*; *MvALS1&3*, populations carrying resistance-conferring mutations in *MvALS1* and *MvALS3*; N.D., populations with no resistance-conferring mutations in *ALS* genes. (c) Methods for resistance diagnosis. In rooting assay, roots were cut at 1 cm from the base of shoots and placed in bensulfuron-methyl solution for 7-10 days. Plants with emerging roots were diagnosed as resistant. In the pot assay, seeds were germinated in a tube, followed by transplantation to pots filled with autoclaved soil. The recommended dose of imazosulfuron (90 g a.i.·ha^-1^) was applied at the three-leaf stage. Plant survival was evaluated at 3 weeks post application. DNA extracted from the plants subjected to sensitivity assays were used for *ALS* gene sequencing.

### Resistance diagnosis in *M. vaginalis*

Resistance to ALS herbicides was diagnosed using the rooting assay with 45 μg·L^-1^ of bensulfuron-methyl, as described by Hamamura et al. (2003). In some populations, the selfed seeds were used for resistance diagnosis using a recommended dose of imazosulfuron [90 g active ingredient (a.i.)·ha^-1^], as described previously (Iwakami et al., 2020). Briefly, seeds were germinated in distilled water in a 1.5 mL tube at 25°C in a climate chamber with a 12 h photoperiod. The germinated seeds were transplanted into autoclaved soil under flooded conditions. When plants reached the three-leaf stage, a commercial formulation of imazosulfuron (75% a.i.) was applied to the surface of standing water. The survival rate of plants was evaluated at 3 weeks post application.

### Imazosulfuron dose–response analysis in *M. vaginalis*

KST1, KOM1, SMM23, AKT3, NRS2, and SMM13 populations were used in the assay. The dose–response analysis was performed as described by Iwakami et al. (2020), with some modifications. Briefly, the seeds were germinated as above, and five seedlings were transplanted into autoclaved soil in 100 cm^2^ plastic pots. The plants were cultivated under flooded conditions. At the two- to three-leaf stage, the commercial formulation of imazosulfuron was applied to the surface of standing water in individual pots at a rate calculated for the area of each pot (0, 0.09, 0.9, 9, 90, 900, 9,000, and 90,000 g a.i.·ha^-1^). At 3 weeks post application, the shoots were harvested and dry weight was measured. The dose that inhibited growth to 50% (GR_50_) and 90% (GR_90_) compared with the control was calculated by fitting a log-logistic curve (two-parameter model) using the drc package (ver. 3.0.1) (Ritz, Baty, Streibig, & Gerhard, 2015).

### Sequencing of *ALS* genes in *M. vaginalis*

DNA was extracted from the leaves of plants subjected to the herbicide sensitivity assay using the cetyltrimethylammonium bromide (CTAB) method (Murray & Thompson, 1980). For some old populations, DNA was extracted from dead seeds using the Plant Genomic DNA Kit (Tiangen, Beijing, China). The entire coding sequence of *MvALS1* to *MvALS5* was PCR-amplified as described by Iwakami et al. (2020). The amplicons were treated with ExoSAP-IT (Affymetrix, CA, USA) to remove dNTPs and primers, and sent for Sanger sequencing to Eurofins Genomics (Tokyo, Japan).

To evaluate the genetic diversity of the whole coding sequence in each *ALS* gene, the nucleotide diversity of each *ALS* gene was calculated using DnaSP 6 (Rozas et al., 2017). The nucleotide diversity at each nucleotide position was also computed to evaluate the difference of the diversity patterns among *ALS* genes. Previously reported full-length coding sequences of three populations (SMM13, SMM23, and TOT1) were also used in the analysis (Iwakami et al., 2020).

### Expression and *in vitro* assay of ALS

The pGEX6P-1 vector (Cytiva, Tokyo, Japan) was digested with SalI-HF (New England Biolabs, Tokyo, Japan). Alleles of *MvALS1/2/3/5* from the sensitive population (SMM13) were individually ligated at the SalI site of pGEX-6 via SLiCE reaction following the protocol described by Motohashi (2015). Each *ALS* gene was expressed without the estimated chloroplast transit peptide (CTP) region corresponding to 1-86 residues of *Arabidopsis thaliana* ALS (Chang & Duggleby, 1997; Singh, Schmitt, Lillis, Hand, & Misra, 1991). The primers used for the SLiCE reactions are listed in Table S2. The resultant expression vector was transformed into *Escherichia coli* BL21 (DE3) cells (Sigma-Aldrich, MO, USA).

The transformed cells were inoculated in 200 mL of LB medium containing 100 μg·ml^-1^ of ampicillin and cultured at 37°C. When OD_600_ reached 0.4-0.7, isopropyl *β*-D-thiogalactopyranoside was added to the medium at a final concentration of 1 mM, and the culture was continued at 16°C for 24 h. The cells were harvested by centrifugation at 14,000 ×*g* for 10 min. The cells were resuspended in phosphate-buffered saline with 1% Triton X-100, 1 mM phenylmethylsulfonyl fluoride, 5 mM dithiothreitol, and 0.5 mM MgCl_2_ and sonicated for 5 min on ice. GST-fused ALS was batch-purified with glutathione Sepharose 4B (Cytiva) according to the manufacturer’ s protocol. ALS was eluted in 400 μL of 50 mM Tris–HCl buffer (pH 8.0) containing 10 mM glutathione, followed by the addition of glycerol at a final concentration of 10% (v/v).

ALS activity was evaluated as described by Chang and Duggleby (1997), with some modifications. The reaction mixture (300 μL) containing 20 mM K_2_HPO_4_/KH_2_PO_4_ (pH 7.0), 50 mM sodium pyruvate, 10 mM MgCl_2_, 1 mM thiamine diphosphate, 20 μM FAD, 0.5 ng enzyme, and varying concentrations of sodium pyruvate (0, 0.78, 1.56, 3.12, 6.25, 12.5, and 25 mM) was incubated at 30°C for 60 min, and the reaction was terminated by adding 30 μL of 6N H_2_SO_4_. The resultant solution was incubated at 60°C for 15 min, followed by the addition of 300 μL creatine (0.5%, w/v) and α-naphthol (0.5%, w/v, in 2.5N NaOH). Absorbance was measured at 525 nm. The data were plotted and curve-fitted to the Michaelis–Menten equation using the drc package (ver. 3.0.1) (Ritz et al., 2015). The Michaelis–Menten constant (*K_m_*) and turnover number (*k_cat_*) was determined for each ALS. The experiments were conducted in triplicate and repeated twice.

Protein concentration was determined using Bio-Rad Protein Assay Dey Reagent Concentrate (Bio-Rad, CA) with bovine serum albumin as the standard, following manufacturer’s instructions.

### Targeted mutagenesis of *ALS* genes

Alleles of *MvALS1*/2/3/5 from the SMM13 population and a deleted allele of *MvALS5* from the SMM3 population were cloned into the pCRblunt vector (Thermo Fisher Scientific, Tokyo, Japan). The codons for Pro197 of the *ALS* genes were mutated to carry a resistance-conferring Ser codon. Targeted mutagenesis was performed according to the protocol in QuikChange Site-Directed Mutagenesis System (Stratagene, CA, USA), with some modifications. Briefly, the clones were used as a template for PCR using PrimeSTAR MAX (TaKaRa, Kusatsu, Japan); the primers are listed in Table S2. The PCR products were treated with DpnI-HF (New England Biolabs). The resultant solution was used for *E. coli* DH5α transformation (ECOS Competent *E. coli* DH5 α, Nippon Gene, Tokyo, Japan) according to the procedures described in the manual. Plasmids were extracted from a culture solution inoculated with a single colony of transformed *E. coli*. The success of the targeted mutagenesis was confirmed by sequencing the extracted plasmids.

### Transformation and phenotyping of transgenic *A. thaliana*

The Pro197Ser-mutated *ALS* genes and the native form of *MvALS1* gene were inserted into the pCAMBIA1390 vector under the cauliflower mosaic virus 35S promoter using the In-Fusion DH Cloning Kit (TaKaRa), as described previously (Iwakami et al., 2019). The primers used in this reaction are listed in Table S2. *A. thaliana* transformation was performed using the agrobacterium-mediated floral dip method as described previously (Guo et al., 2019). The transformants were selected with hygromycin B (20 mg· L^-1^).

To quantify transgene expression, real-time RT-PCR was performed. Briefly, RNA from 10-day-old T3 homozygous lines was extracted using the Plant Total RNA Mini Kit (Favorgen, Ping-Tung, Taiwan). Contaminated DNA in the extracted RNA was removed using the Turbo *DNA-free* kit (Thermo Fisher Scientific). The resultant RNA was reverse transcribed according to the method described by Iwakami et al. (2019). PCR was run using THUNDERBIRD SYBR qPCR Mix (TOYOBO, Osaka, Japan) on QuantStudi 12K Flex Real-Time PCR System (Thermo Fisher Scientific). The primers for transgene expression were designed using the terminator sequence of pCAMBIA1390 (Table S2). *GAPDH* was used as the internal control; primers developed by Czechowski, Stitt, Altmann, Udvardi, and Scheible (2005) were used. Data were analysed using the ΔΔCT method (Schmittgen & Livak, 2008).

Imazosulfuron sensitivity of the transformants was evaluated as described previously (Dimaano, Yamaguchi, Fukunishi, Tominaga, & Iwakami, 2020) with some modifications. Seeds were sown on Murashige and Skoog medium (Murashige & Skoog, 1962) containing various concentrations of imazosulfuron. Plants were grown for 10 days at 22°C. Sensitivity was evaluated based on the emergence of true leaves. GR_50_ was computed as described above.

### Gene expression analysis in *M. vaginalis*

SMM13 and SMM23 were used as representatives of the sensitive and resistant populations, respectively. Germinated seeds were transplanted into autoclaved soils. Six plants from each population were grown in water until the three-leaf stage. Half of the plants were treated with imazosulfuron at a concentration of 90 g a.i.·ha^-1^. At 24 h post application, the shoots of all plants were snap frozen in liquid nitrogen and stored at −80°C until further use. RNA was extracted using the RNeasy Plant Mini Kit (Qiagen, Venlo, Netherlands), followed by DNase treatment using the Turbo *DNA-free* kit (Thermo Fisher Scientific).

To estimate approximate transcript abundance, PCR products generated by the primers designed based on the completely conserved regions among the five copies of *ALS* genes (Table S2) were cloned into the pCRblunt vector (Thermo Fisher Scientific). Over 50 clones were sent to Eurofins Genomics for sequencing. cDNA was prepared from the RNA of an imazosulfuron-treated sensitive plant. The experiment was performed twice.

For RNA-Seq analysis, 12 RNA samples were shipped to GENEWIZ (Kawaguchi, Japan) for stranded RNA-Seq using Illumina HiSeq 4000 (150-bp paired-end reads). Contaminated adaptor sequences were trimmed using ‘cutadapt’ (ver. 2.5) (Martin, 2011). Low-quality bases were removed using Trimmomatic (ver. 0.38) (Bolger, Lohse, & Usadel, 2014) with the following options: ‘LEADING:25 TRAILING:25 SLIDINGWINDOW:4:15 MINLEN:100’. Reads of a non-treated sensitive plant were used for contig assembly. The assembly was performed using Trinity (ver. 2.4.0) (Grabherr et al., 2011) with a k-mer size of 32. The biological function of the contigs was inferred using BLASTX (Blast+ ver. 2.9.0) (Camacho et al., 2009) against rice protein sequences (*Oryza sativa* MSU release 7). Only one contig under the identical gene identifier annotated by Trinity was used, as described by Ono et al. (2015). Briefly, from contigs with hits in the BLAST analysis, those with the highest bit score were selected, and from contigs without hits, the longest ones were selected. Contigs that were annotated as *ALS* genes were replaced with full-length *ALS* genes of *M. vaginalis* (DDBJ accession numbers: LC488950 to LC488953 and LC488956). The reads were mapped using Bowtie2 (ver. 2.3.4.1) (Langmead & Salzberg, 2012) with the following options: ‘-X 900 --very-sensitive --no-discordant --no-mixed --dpad 0 --gbar 99999999’. The concordantly mapped reads were counted using RSEM (ver. 1.3.2) (Li & Dewey, 2011). Multiple comparisons were performed using the GLM approach with the quasi-likelihood F-test using the edgeR package (ver. 3.28.1.) (Robinson, McCarthy, & Smyth, 2010). Contigs with fold-changes of more or less than 2 and *q*-values less than 0.05 were evaluated as differentially expressed genes (DEGs).

## RESULTS

### Herbicide resistance of *M. vaginalis* collected across Japan

In this study, we sampled 113 suspected resistant populations across Japan and tested for ALS herbicide sensitivity using the rooting or pot assays (Figure 1). Sixty-six populations (58.4%) were confirmed to be resistant to ALS herbicides. All the resistant populations carried one or two homozygous mutations in the codons of *ALS* genes that are known to confer resistance to the ALS herbicides, namely Pro197, Ala205, and Asp376, implying that *M. vaginalis* tend to evolve resistance to ALS inhibitors via target-site resistance (TSR) mechanisms as reported in many non-Poaceae weeds (Yu & Powles, 2014).

Analysis of the codons involved in ALS resistance in the 66 populations tested in this study and 2 previously reported resistant populations (Iwakami et al., 2020) revealed 9 types of resistance-conferring AASs. Intriguingly, the identified resistance-conferring mutations were exclusively found in *MvALS1* and *MvALS3*, and not in other three genes (Figure 2a). Mutations at Pro197 dominated the others (Figure 2b), probably because sulfonylureas, a chemical class known to select for Pro197 mutation (Yu & Powles, 2014), have been the major ALS herbicides used in Japan (Hamamura, 2018). There was no clear association between mutation type (i.e. AASs and genes) and geographical region (Figure 1b; Table S1). As relatively rare cases, populations with two independent mutations were also found. AIZ2 and KR1 populations carried two resistance-conferring mutations, namely Pro197Ser and Asp376Glu, in a single gene (*MvALS1*), while YNZ1 population carried Asp376Glu in *MvALS1* and Pro197Ser in *MvALS3*. In addition, mutations that have not been reported in natural weed populations, including Asp376Asn in NDA1 and Ala205Thr in ANP1, were also detected. Both mutations were confirmed to confer reduced sensitivity to sulfonylurea herbicides in yeast (Bedbrook et al., 1995; Duggleby, Pang, Yu, & Guddat, 2003). Thus, AASs may be the causal mutations for resistance in these weed populations, although it should be noted that we did not actively screen for additional non-target site mechanisms.

**Figure 2.**
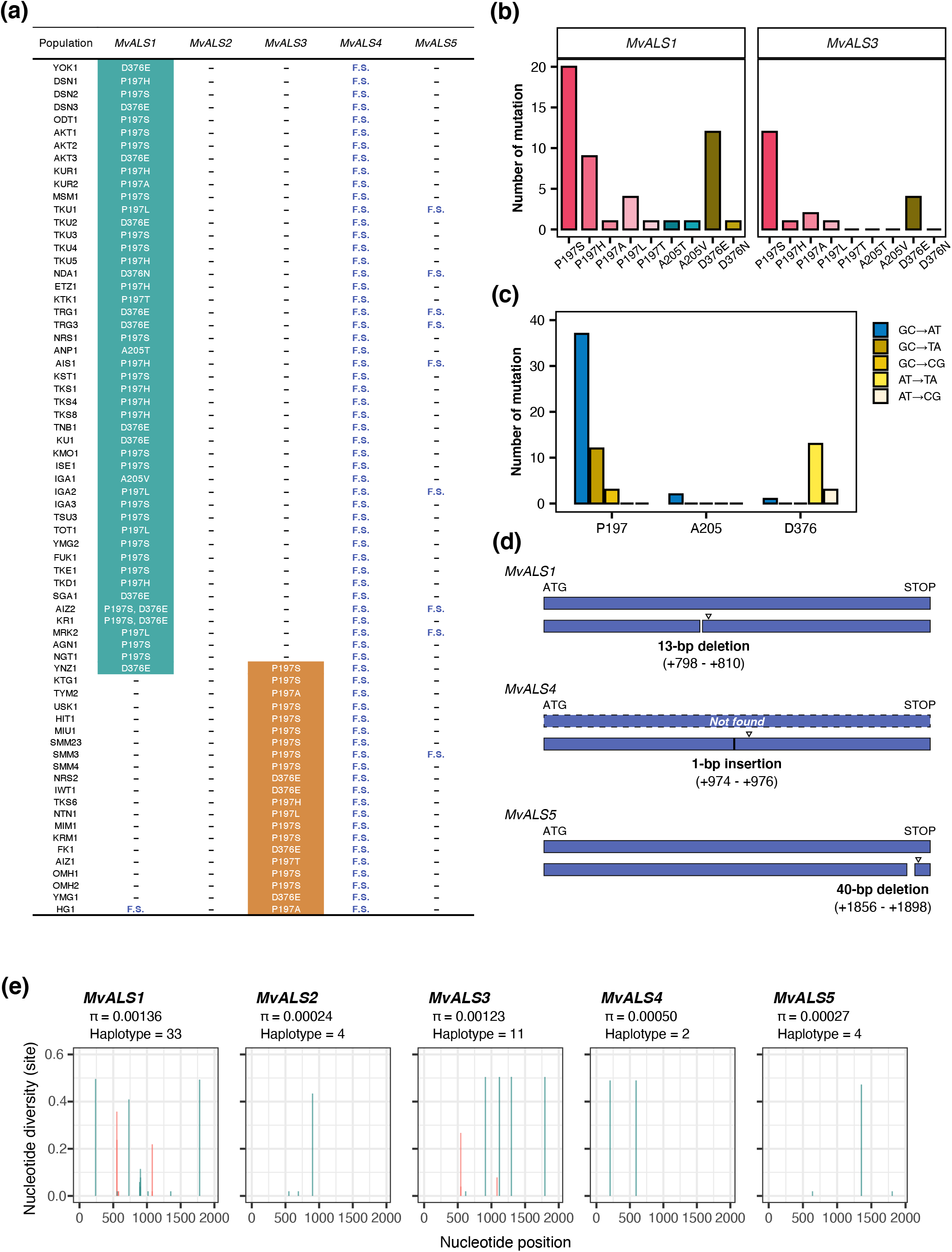
*ALS* genes of *Monochoria vaginalis*. (a) Status of *ALS* genes in 68 resistant populations analysed in this study. YNZ1 carries mutations both in *MvALS1* and *MvALS3*. AIZ2 and KR1 populations carry two independent mutations in *MvALS1*. F.S., frameshift mutation. (b) Patterns of resistance-conferring mutations. (c) Patterns of DNA conversions of resistance-conferring mutations. (d) Schematic representation of frameshifted alleles found in *MvALS1, MvASL4*, and *MvALS5. MvALS4* alleles encoding the canonical form of ALS were not detected in the present study. Triangles represent a premature stop codon. (e) Nucleotide variations in the five *ALS* genes. The coding sequences of 68 resistant and 31 sensitive populations were analysed. The plot shows nucleotide diversity calculated at each nucleotide position. Positions causing an amino acid replacement that confers resistance are indicated in red. Nucleotide diversity of the entire genes (π) and haplotype numbers were calculated.

Transition was the dominant type of resistance-conferring mutation (Figure 2c), consistent with the results of a previous genome-wide mutation analysis of *A. thaliana* (Weng et al., 2019). The highest frequency of the Pro197Ser transition mutation (CCT→<u>T</u>CT) may be partly attributed to the nature of mutation patterns, although other factors such as high level resistance caused by this mutation (Sada, Ikeda, & Kizawa, 2013) may play a more important role in the observed bias.

To gain insights into the mystery of the exclusive presence of the TSR allele in only two *ALS* loci, we further analysed the coding sequence of each *ALS* gene in 66 resistant and 30 sensitive populations, along with the previously reported sequences of 2 resistant and 1 sensitive populations (Iwakami et al., 2020) (Table S1). Coding sequences of all *ALS* genes in *M. vaginalis* were homozygous, contrary to the observation in obligate outcrossing weeds such as *Lolium rigidum* and *Papaver rhoeas* (Délye, Boucansaud, Pernin, & Le Corre, 2009; Délye, Pernin, & Scarabel, 2011). This result highlights the predominant self-pollinating nature of *M. vaginalis*. Consistent with previous reports (Iwakami et al., 2020; Ohsako & Tominaga, 2007), a T insertion causing a frameshift was detected in all *MvALS4* alleles investigated in this study (Figure 2a-d). The products encoded by these mutant alleles lacked domains with pivotal roles in ALS activity, indicating that this gene does not encode the functional form of ALS. Thus, the prevalence of frame-shifted alleles in *MvALS4* among *M. vaginalis* populations in Japan explains the absence of resistance-conferring mutations in this gene. While all alleles were pseudogenised in *MvALS4*, variations were observed in *MvALS1* and *MvALS5*; as such a 13 bp deletion was detected in *MvALS1* in 1 population and a 40 bp deletion in *Mv*ALS5 in 12 populations (Figure 2a, TableS1). The reading frame of these alleles also shifted, resulting in premature stop codons (Figure 2d). The putative loss-of-function alleles of *MvALS5* were found in populations sampled from various areas without any geographical association. Contrary to that in *MvALS4*, majority of the alleles were not pseudogenised in *MvALS1* and *MvALS5*.

The analysis of coding sequences revealed that genes with resistance-conferring alleles (*MvALS1* and *MvALS3*) showed a higher nucleotide diversity and greater variation in haplotype number (Figure 2e). This trend held true even when naturally selected resistance-conferring alleles were excluded from the analysis: a greater nucleotide diversity (per site) was observed at three and four positions, irrespective of the resistance selection, in *MvALS1* and *MvALS3*, respectively. Thus, *MvALS1* and *MvALS3* might be more prone to spontaneous mutations, some of which may occur by chance at the resistance-conferring codons. However, the difference in the rate of spontaneous mutations among genes can hardly explain the all-or-nothing resistance-conferring alleles of the *ALS* gene family in *M. vaginalis*. Other more pronounced factors may have played key roles in shaping these biased patterns.

### Functional characterisation of each ALS

To elucidate the mechanism underlying the disproportionate distribution of TSR mutations among *ALS* copies, we next compared the enzymatic functions of the protein encoded by the canonical alleles of each *ALS* gene. ALS activity of the purified GST-tagged protein was assessed (Figure S1). The activity of the recombinant proteins was evaluated based on the conversion of the produced acetolactate to a red insoluble complex. The reaction solutions of the recombinant proteins of MvALS1/2/3/5 developed a red colour, indicating that all isozymes produced acetolactate (Figure 3a). The enzymatic properties of the respective isozymes were similar, although the affinity to the substrate pyruvate (*kcat/Km*) was relatively high in MvALS1.

**Figure 3.**
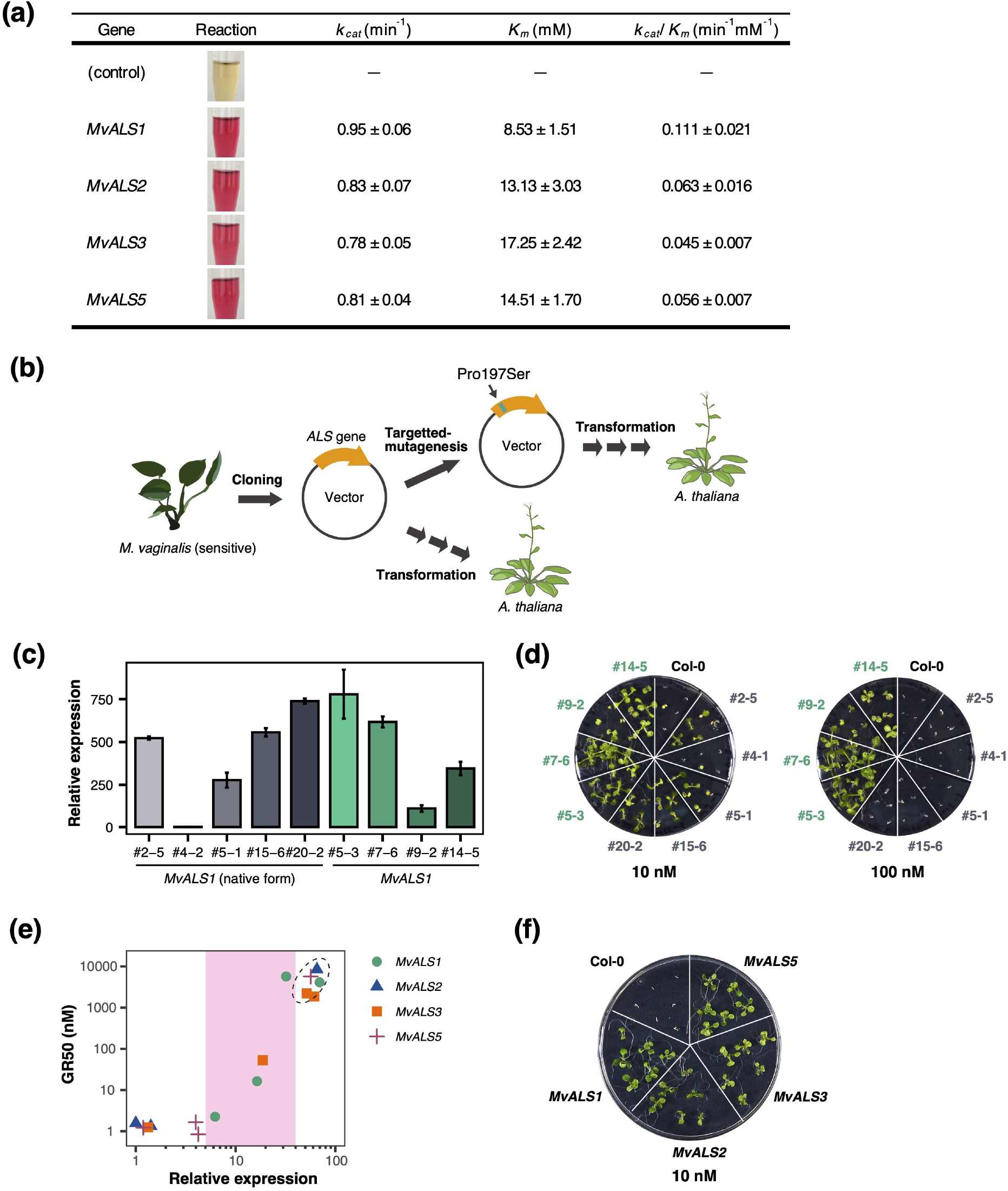
Enzymatic functions of ALS encoded by each gene of *Monochoria vaginalis*. (a) Enzymatic properties of recombinant ALS. (b) Transformation of *Arabidopsis thaliana. ALS* genes were isolated from a sensitive plant, and were mutagenised to encode the Pro197Ser form of ALS. The native form of *MvALS1* was also transformed. (c) *MvALS1* expression in transgenic *A. thaliana* lines. Bars represent standard error (n=3). (d) Response of transgenic *A. thaliana* to 10 and 100 nM imazosulfuron. (e) Relationship of transcript level and imazosulfuron sensitivity in transgenic *Arabidopsis thaliana* expressing each *ALS* gene with Ser at Pro197 codon. (f) Response of transgenic *A. thaliana* to 10 nM imazosulfuron. The plants correspond to the lines indicated by circles in Figure 3b.

We examined the potential of TSR mutation to lead to herbicide resistance in all loci under consideration by transforming *A. thaliana*. In this study, we artificially generated Pro197Ser alleles (Figure 3b), which are known to endow extreme resistance to some ALS herbicides (such as imazosulfuron) in other species (Sada et al., 2013). First, we investigated the impact of the TSR mutation by comparing the imazosulfuron sensitivity of *A. thaliana* lines carrying the *MvALS1* gene with or without the TSR mutation. We generated five and four single-copy lines of the native allele and the TSR allele with varying transcript levels (Figure 3c). Lines with the TSR allele survived the lethal dose of 10 nM, and even 100 nM (Figure 3d). On the other hand, lines with the wild-type allele exhibited only a marginal resistance level, with plants in the highest expression lines dead at 100 nM. The results indicate that the qualitative, not quantitative, alteration in the ALS target site plays a key role in the evolution of resistance to ALS herbicides.

Next, we investigated the sensitivity of *A. thaliana* lines with the TSR allele of the respective *MvALS* gene. Interestingly, all the TSR alleles conferred imazosulfuron resistance in *A. thaliana* when overexpressed (Figure 3e-f). Resistance indices of all transgenic lines were correlated with the transcript level of transgenes (*ALS* genes of *M. vaginalis*), and a similar resistance level was observed when the transcripts of the respective *ALS* genes were accumulated at a similar level. Notably, even a small difference in transcript level drastically affected herbicide sensitivity when the transcript levels were within a specific range. For instance, although the transcript levels in three *MvALS1* transgenic lines (lines indicated in pink area in Figure 3b) differed by only 5.1-fold at the maximum, their resistance levels differed by as much as 2,500-fold. Meanwhile, very low or high transcript levels did not affect sensitivity.

We also tested the putative loss-of-function allele of *MvALS5*, as it only partially lacks the γ domain—one of the three ALS domains (Figure 2d). All T2 generations in which the allele of *MvALS5* carrying the Pro197Ser mutation was deleted were sensitive to imazosulfuron, while at least seven lines carrying the canonical allele of *MvALS5* showed decreased imazosulfuron sensitivity (Figure S2). This result implies that the deleted form of *MvALS5* does not encode functional ALS.

Collectively, the results of recombinant protein assay and ectopic expression in *A. thaliana* suggest that the activities of the four isozymes were near identical, although a slightly higher substrate specificity was observed in MvALS1 in the recombinant assay. Notably, the recombinant ALS assay is affected by multiple factors, such as the prediction of the CTP cleavage site and presence of a GST tag, which may have influenced the results for MvALS1. Our data suggest that *MvALS2* and *MvALS5* could potentially be involved in the evolution of ALS resistance in *M. vaginalis* contrary to the expectation from the nation-wide survey (Figure 2).

### Transcript levels of *MvALS* genes

Since no significant differences were observed in recombinant protein assays and ectopic expression in *A. thaliana*, we compared the transcript levels of *ALS* genes in *M. vaginalis* seedlings in the presence or absence of imazosulfuron using mRNA-Seq. *De novo* assembly followed by the curations of *ALS* contigs, we obtained 84,649 contigs with average length of 802.5 bp.

The transcriptome of herbicide-treated sensitive plants formed a distinct cluster (Figure 4a), indicating that herbicide treatment triggered massive reprogramming of gene expression within 24 h in these plants. Statistical analysis revealed 12,845 differentially expressed contigs between the treated and untreated sensitive populations (Figure S3). In contrast, gene expression in the resistant population remained almost unchanged following herbicide treatment, which was expected based on a previous report that herbicide treatment only slightly suppressed growth in this resistant population (Iwakami et al., 2020). Moreover, the difference between the untreated sensitive and resistant populations was minor, with only 528 differentially expressed contigs (Figure S3).

**Figure 4.**
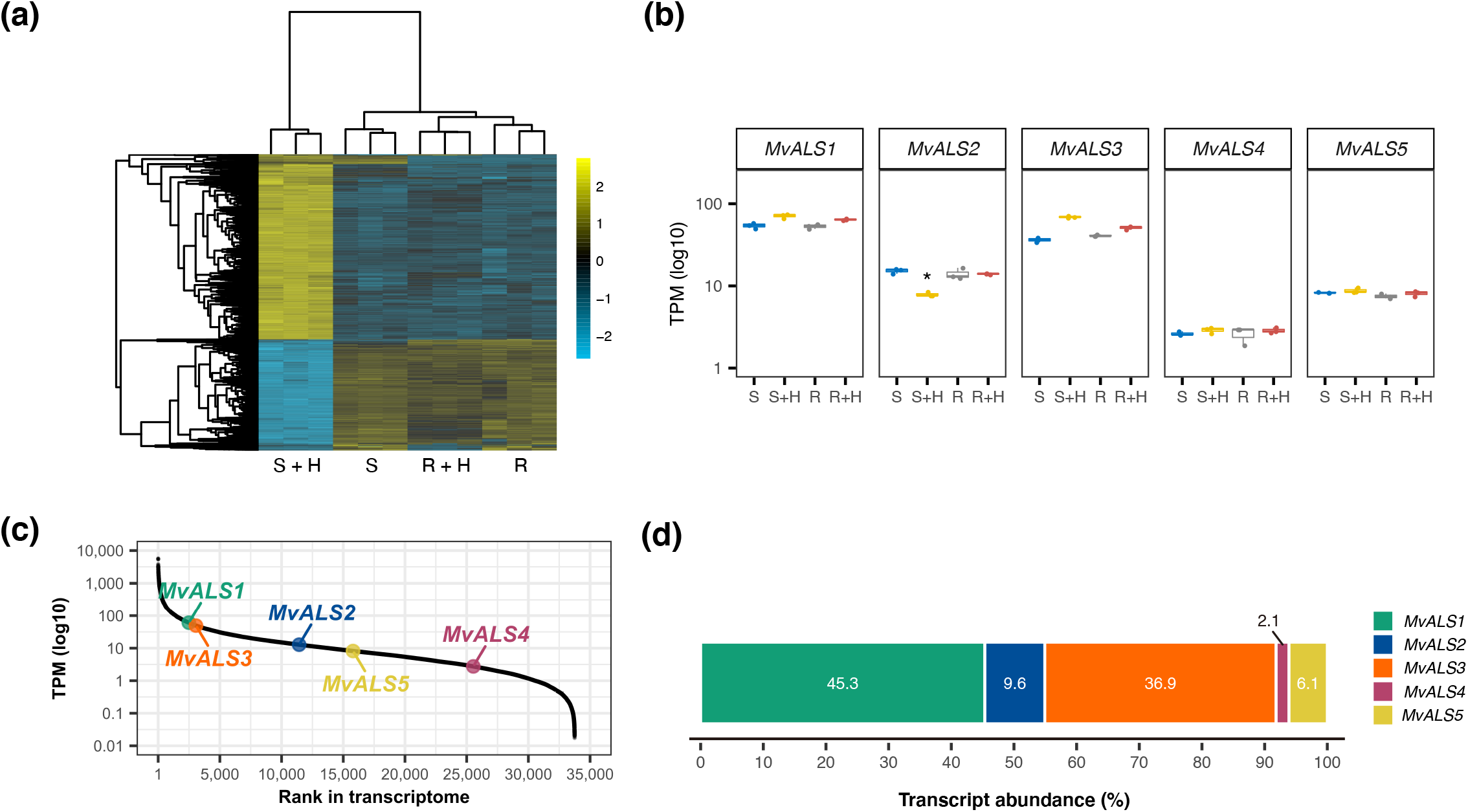
Expression of *ALS* genes in *Monochoria vaginalis*. (a) Transcriptome of sensitive (S) and resistant (R) *M. vaginalis* with or without herbicide imazosulfuruon (H) application. Transcriptome of the R plants was almost unchanged, while that of the S plants was drastically affected. (b) Transcripts per kilobase million (TPM) of each *ALS* gene. \**q*<0.05. (c) Expression rank in *M. vaginalis* transcriptome. Data represent the average of 12 mRNA-Seq samples. (d) Relative transcript abundance of *ALS* genes in *M. vaginalis*.

The transcription of *ALS* genes was almost stable among all the samples (Figure 4b). The only exception was *MvALS2*, which was slightly downregulated (2.36-fold, *q*<0.01) in herbicide-treated sensitive plants. As the differences among the populations were small, we averaged the transcripts per kilobase million (TPM) data of 12 libraries and examined the transcript level of each *ALS* gene. The TPM value of *MvALS1* and *MvALS3* ranked 2,487 and 3,062, respectively, and these values were much higher than the TPM values of *MvALS2, MvALS5*, and *MvALS4* (Figure 4c-d). Moreover, respectively 45.3% and 36.9% of the reads mapped to the *ALS* genes were from *MvALS1* and *MvALS3*, followed by 9.6% from *MvALS2*. These findings were consistent with the results of gene cloning (Figure S4). Therefore, the two genes carrying resistance-conferring mutations (*MvALS1* and *MvALS3*) in natural *M. vaginalis* populations were among the most highly expressed *ALS* genes. The absence of a significant difference in the transcript levels of each copy between the sensitive and resistant populations under non-treated conditions implies that the observed diversity of transcription among *ALS* genes is not an evolutionary response to the selection of ALS herbicides.

### Roles of *MvALS1* and *MvALS3* in resistance evolution

We compared the roles of *MvALS1* and *MvALS3* in the resistant phenotype of *M. vaginalis*. Imazosulfuron sensitivity of populations carrying the same mutation in *MvALS1* or *MvALS3* was compared. Three populations with the Pro197Ser mutation showed similar GR_50_ and GR_90_ values, irrespective of the mutated genes (Figure 5). Moreover, similar GR_50_ and GR_90_ values were observed in another set of populations carrying the Asp376Glu mutation. Therefore, the mutated alleles of *MvALS1* or *MvALS3* equally contribute to resistance in *M. vaginalis*.

**Figure 5.**
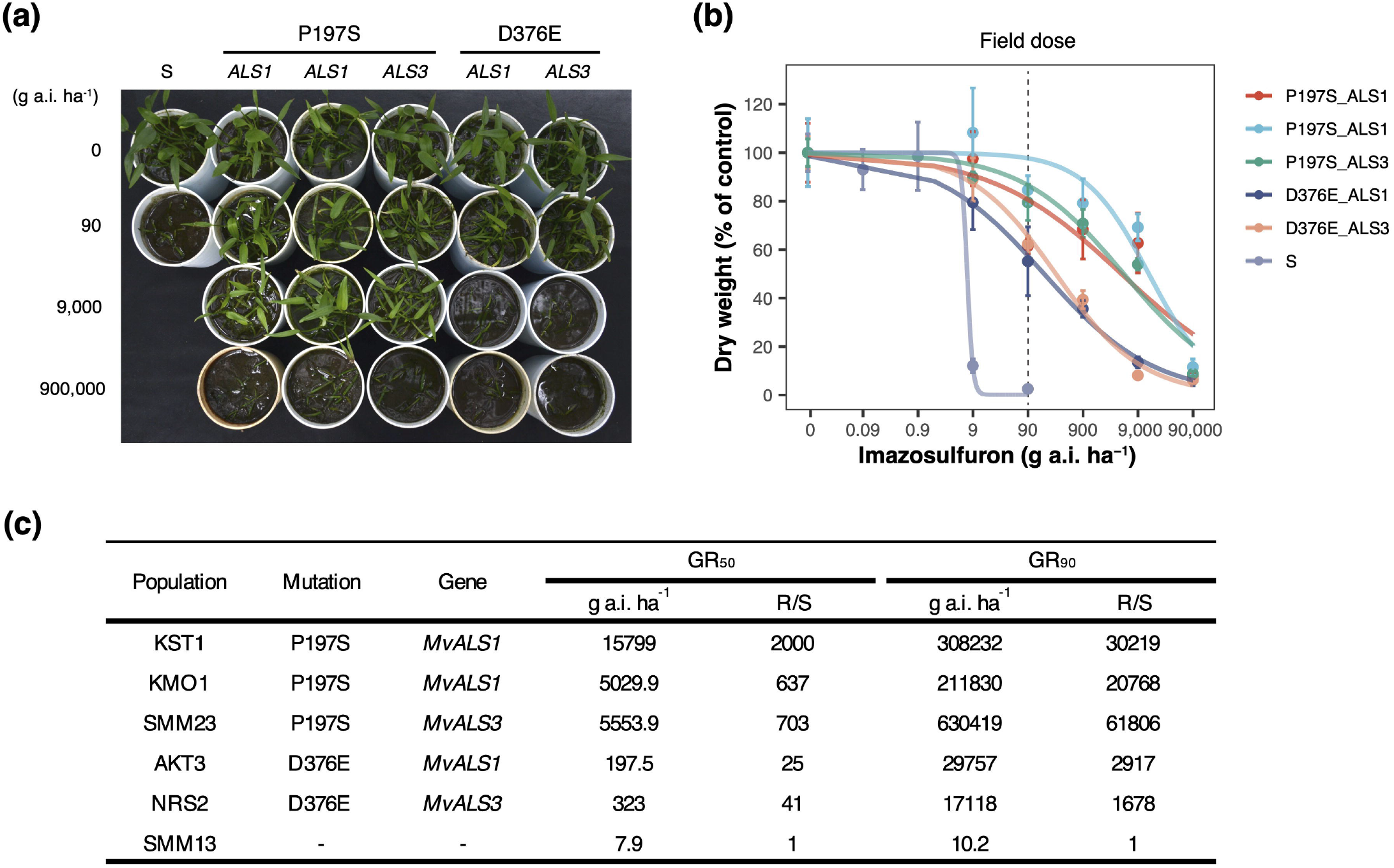
Herbicide sensitivity of *Monochoria vaginalis* carrying a mutation in *MvALS1* and *MvALS3*. (a) *M. vaginalis* 3 weeks post imazosulfuron application. (b) Dose response to imazosulfuron. Bars represent standard error (n=3). (c) Doses that inhibited the growth to 50% (GR_50_) and 90% (GR_90_).

## DISCUSSION

Discerning the evolutionary genetic paths available for adaptation to a particular environment opens the door to predicting evolutionary dynamics and controlling the evolution of the species. In the evolution of weed resistance to herbicides, evolutionary paths are often restricted. In particular, in resistance to ALS herbicides in broadleaf weeds, extreme parallelism in *ALS* genes is often observed, implying that almost no genetic solutions are available to adapt to the herbicides except for the alteration of *ALS* genes. Meanwhile, it remains unclear whether all *ALS* loci represent potential evolutionary paths for herbicide resistance when multiple copies of the *ALS* gene exist in the species of interest. To investigate whether duplicated genes generate additional chances for resistance evolution, we studied the molecular mechanisms underlying the evolution of herbicide resistance in the noxious paddy weed *M. vaginalis*, taking advantage of the highest copy number of *ALS* genes reported so far in a weed. Through the study of >100 populations, we revealed that the Japanese populations of *M. vaginalis* utilised only two *ALS* loci, *MvALS1* and *MvALS3*, for TSR evolution among the five *ALS* loci in the genome. The subsequent molecular characterisations of each *ALS* gene strongly suggest that the exclusivity was underpinned by the large difference in the expression of each gene: the transcripts of *MvALS1* and *MvALS*3 are more abundant than those of the other *ALS* genes. This study revealed that among the multiple potential hotspots in the genome of *M. vaginalis*, expression level confers evolutionary competency in two *ALS* loci, shaping the biased pattern of resistance-conferring alleles in resistant populations of *M. vaginalis* in Japan.

The generation of evolutionary innovations derived from gene duplication is often addressed in evolutionary biology (Panchy, Lehti-Shiu, & Shiu, 2016; Xu, Wang, Guo, He, & Shi, 2020). While attention has been paid to the acquisition of novel functions or increased protein abundance by duplicate genes, little is known about their roles in the expansion of evolutionary paths. When the molecular function of the encoded protein is conserved, all duplicates can be considered to have potential for evolutionary innovation. In fact, in weeds with multiple copies of *ALS* genes such as *Schoenoplectiella juncoides* (Sada & Uchino, 2017), *Descurainia sophia* (Xu, Xu, Li, & Zheng, 2020), *Senecio valgaris* (Délye et al., 2016), and *Echinochloa crus-galli* (Löbmann, Schulte, Runge, Christen, & Petersen, 2021), all the loci have been used for resistance evolution, suggesting that the duplicate copies increase the chances of resistance evolution. On the other hand, this study revealed that the independent selective pressure exerted in many Japanese agricultural fields exclusively selected only two *ALS* loci for evolutionary alteration, suggesting that duplicated loci do not necessarily generate available evolutionary tools, at least in the context of microevolution, even if the molecular function of the encoded protein is conserved (see discussion below).

Functional characterisation of *MvALS1/2/3/5* revealed no marked differences in their functions, indicating that *MvALS2* and *MvALS5*, which did not carry resistance-conferring non-synonymous mutations in natural populations, are the ‘potential’ hotspots for resistance evolution, similar to *MvALS1* and *MvALS3*. However, since the two loci are transcriptionally not very active in seedlings, resistance-conferring mutations in these *ALS* genes would not confer herbicide resistance in agricultural fields under the current expression profiles of the *ALS* genes in *M. vaginalis*. Assuming that the transcript abundance of *ALS* genes is proportional to the amount of ALS protein, MvALS2 and MvALS5 accounted for <10% of the ALS pool, whereas MvALS1 and MvALS3 accounted for nearly 40%. This result indicates that over 90% ALS would be inhibited by herbicides if a resistance-conferring mutation is present in *MvALS2* or *MvALS5;* this proportion would further decrease in the first generation of plants carrying a spontaneous mutation due to the heterozygous status, and such plants would likely not survive herbicide stress in agricultural fields. In addition to relative abundance, absolute transcript abundance may also play important roles. We observed that *MvALS1* and *MvALS3* were among the most transcriptionally active genes in *M. vaginalis*, positioned close to the first inflexion point on the sigmoid curve of the transcription ranking plot, while the other three genes were in the middle or lower parts of the curve (Figure 4c). Interestingly, three *ALS* genes in hexaploid wheat, each of which confers resistance when Pro197 is substituted via targeted mutagenesis (Zhang et al., 2019), were also placed close to the first inflexion point (Figure S5a). These observations, together with the mRNA–GR_50_ relationships in transformed *A. thaliana* (Figure 3b), suggest that a certain degree of transcript abundance may be required for protein expression. Thus, in order for *MvALS2* and *MvALS5* to be involved in resistance, an exceptionally large increase in gene expression or two sequential mutations, a gene up-regulation followed by a non-synonymous substitution, would be required. The impact of the former, as evident from the *A. thaliana* transformation experiment (Figure 3c-d), would be only marginal, which would hardly endow appreciable resistance in the fields, considering that a >10 times higher resistance level is required for wild-type *M. vaginalis* to tolerate the field dose of imazosulfuron (90 g a.i.·ha^-1^) (Figure 5b). The latter is theoretically possible, but would require two independent mutations, one qualitative and one quantitative, in the same individual, implying that this rarely occurs. The most likely scenario of *MvALS2/5-based* TSR evolution would be a non-synonymous TSR substitution in populations with a defect in a highly expressed *ALS* gene, such as the HG1 population (Figure 2a), where the relative dominance of *MvALS2/5* to the whole ALS pool is higher. Currently, individuals with loss-of-function alleles of *MvALS1* are extremely rare in Japan. If this genotype increases in the future, individuals with TSR mutations in *MvALS2/5* may arise. It will be important to study the effects of *MvALS1* deficiency on fitness to estimate the evolutionary dynamics of this genotype.

Although our data strongly underline the role of gene expression in the biased representation of mutated genes inducing resistance, we cannot rule out the role of other factors. Firstly, given the observed higher sequence variations in *MvALS1* and *MvALS3* (Figure 2e), the spontaneous mutation rate might be higher for these genes, leading to their overrepresentation. According to the classic evolutionary theory, the spontaneous mutation rate is random across a genome, and the apparent bias is the consequence of fitness costs. However, a recent sequencing study revealed that cytogenetic features influenced the mutation rate itself to a certain degree (Monroe et al., 2020). Thus, the spontaneous mutation rate may drive the biased representation of genes. Secondly, multi-layered DNA repair systems (Kimura & Sakaguchi, 2006) may also play a role in the bias. These repair mechanisms are affected by the flanking sequences, local DNA structure, and gene expression, among other factors, and do not act uniformly throughout the genome. The local DNA environment of each *ALS* locus may have affected the manifestation of spontaneous DNA mutations. Thirdly, our gene expression results must be interpreted with caution, as we analysed only two populations. There may be natural variations in *ALS* gene expression among *M. vaginalis* populations. However, similar herbicide responses observed among populations carrying the same mutations in this study (Figure 5) may alternatively indicate relatively conserved gene expression among *M. vaginalis* populations in Japan. Large-scale analyses, such as population transcriptomics, combined with comprehensive investigations of differential mutation rates in the *ALS* gene family would provide a more holistic view of evolutionary patterns in *M. vaginalis*.

Notably, resistance-conferring mutations were more frequent in *MvALS1* than in *MvALS3* (Figure 2a). This may be because a mutation in *MvALS1* may confer higher resistance than that in *MvALS3*, which is beneficial to plants, particularly those carrying mutations conferring low resistance, such as Pro197Leu (e.g. Sada et al., 2013), in the heterozygous status. However, comparison of plants carrying the identical mutations in different genes did not reveal significant differences in herbicide sensitivity (Figure 5), which is in agreement with an independent study (K. Ohta, Y. Fujino & Y. Sada, unpublished). Furthermore, in transformed *A. thaliana, MvALS1* and *MvALS3* conferred a similar level of resistance when expressed equally (Figure 3b). Thus, although higher substrate affinity was detected for MvALS1 in our recombinant enzyme assay (Figure 3a), other factors (e.g. the bias of material sampling) were likely responsible for the bias between *MvALS1* and *MvALS3*.

In conclusion, comprehensive analysis of *ALS* gene sequences in Japanese populations of *M. vaginalis* revealed that variable transcript levels of these genes are associated with the biased patterns of mutations for the evolution of herbicide resistance. This study provides a novel insight that gene expression quantitatively provides evolutionary competency to potential hotspots across genome for resistance evolution. In other words, gene expression level can act as a genomic constraint for the parallel evolution of herbicide resistance. Of note, the Japanese populations of *M. vaginalis* may only reflect a particular genetic group of this globally distributed weed (mainly in eastern and southern Asia), considering that Japan is a small island country. Thus, further large-scale studies of the global populations would provide a more comprehensive understanding of the evolutionary strategies of herbicide resistance in *M. vaginalis*, which may not be limited to the two hotspots (*MvALS1* and *MvALS3*) detected in this study. In the era of genomics, it is anticipated that numerous genomes with more complex DNA structures, such as polyploid genomes, will be compiled, which will inevitably impose more attention to the roles of redundant copies during the course of evolution. The perspective and framework of this study are applicable to research on TSR evolution in agricultural weeds as well as broader studies focusing on the process of bias for genetic loci in parallel evolution.

## ACKNOWLEDGEMENTS

We thank Nihon Nohyaku for providing seeds of *Monochoria vaginalis*, and members of the weed science lab in Kyoto University for the cooperation to collect *M. vaginalis*, Yuuri Nakamura for technical assistance. Computations were partially performed on the NIG supercomputer at ROIS National Institute of Genetics. This work was partly supported by Japan Association for Advancement of Phyto-Regulators (JAPR) grant no. 2017-4, 2018-4 (to A.U. and S.I.), 2019-2 (to S.O. and S.I.).

## AUTHOR CONTRIBUTIONS

Project Conception, S.I.; Experimental Design, S.T., T.M., and S.I.; Performance of Experiments, S.T., A.U., S.O., S.I.; Bioinformatics Analysis, S.T. and S.I.; Material Collection and/or Maintenance, S.T., A.U., S.O., C.M., K.H., M.M., N.Y., N.U., Y.T., N.F., E.K., Y.S., T.T., and S.I.; Visualization, S.T. and S.I.; Writing, S.I.; Supervision, T.T., S.I.

## DATA ACCESSIBILITY

mRNA-Seq data have been deposited in the DDBJ Sequence Read Archive (DRA) database with BioProject ID PRJDB11326 (BioSample: SSUB017551).

## Notes

### Competing Interest Statement

The authors have declared no competing interest.

